# Sensory neuron-derived Na_v_1.7 contributes to dorsal horn neuron excitability

**DOI:** 10.1101/551747

**Authors:** Sascha R.A. Alles, Filipe Nascimento, Rafael Luján, Queensta Millet, Ali Bangash, Sonia Santana, James J. Cox, Marco Beato, Jing Zhao, John N. Wood

**Affiliations:** Molecular Nociception Group, Wolfson Institute for Biomedical Research, University College London, London WC1E 6BT, UK; Department of Neuroscience, Physiology and Pharmacology, University College London, London WC1E 6BT, UK; Synaptic Structure Laboratory, Instituto de Investigación en Discapacidades Neurológicas (IDINE), Department Ciencias Médicas, Facultad de Medicina, Universidad Castilla-La Mancha, Campus Biosanitario, C/Almansa 14, 02008 Albacete, Spain

## Abstract

Expression of the voltage-gated sodium channel Nav1.7 in sensory neurons is required for pain sensation. We examined the role of Nav1.7 in the dorsal horn of the spinal cord using an epitope-tagged knock-in mouse. Immuno-electron microscopy showed the presence of Nav1.7 in dendrites of lamina II neurons, despite the absence of mRNA. Peripheral nervous system-specific Nav1.7 KO mice showed central deficits with lamina II dorsal horn tonic firing neurons more than halved and single spiking neurons more than doubled. Nav1.7 blocker PF05089771 diminished excitability in dorsal horn neurons, but had no effect on Nav1.7 KO mice. These data demonstrate an unsuspected functional role of peripherally generated Nav1.7 in dorsal horn neurons and an expression pattern that would not be predicted by transcriptomic analysis.

## Introduction

The problem of pain continues to grow, and new analgesic approaches are urgently required (*1*). Human genetic studies have identified a number of potential analgesic targets including neurotrophins, transcription factors and ion channels (*2-4*). The sodium channel Nav1.7 expressed in sensory neurons is required for pain perception in mice and humans (*5*). Nav1.7 gain-of-function mutations result in ongoing pain in humans, whilst rare recessive loss of function mutants are pain free, but otherwise normal apart from an inability to smell (*6*). Nav1.7 has been assumed to play an essential role in generating nociceptive spiking in sensory neurons. However, it has other roles in pain pathways. In mice, deletion of Nav1.7 in sensory neurons leads to enhanced expression of opioid peptides as well as potentiated opioid receptor activity (*7, 8*). Much of the analgesia associated with loss of function of Nav1.7 in mice (and one human tested loss-of-function patient) can be reversed with the opioid antagonist naloxone, whilst Nav1.7 antagonist action is greatly potentiated by low dose opioids or enkephalinase blockers (*9, 10*). Nav1.7 also has an unusual role as an integrator of synaptic input in the hypothalamus (*11*).

Anosmia associated with loss of Nav1.7 in olfactory neurons has been shown to result from lack of glutamate release (*6*). In somatosensory neurons there is also a loss of substance P release from Nav1.7 null mutant sensory neurons suggesting a regulatory role for Nav1.7 in the control of neurotransmitter release (*12*). To examine this mechanism we generated an epitope tagged Nav1.7 knock in mouse that shows entirely normal pain behaviour. We used this mouse first to identify Nav1.7 interacting proteins using immunoprecipitation and mass spectrometry (*13*) and then to examine the expression of Nav1.7 in the central terminals of sensory neurons using both immunocytochemistry and immuno-electron microscopy.

Immunohistochemical studies showed that the TAP-tagged Na_v_1.7 is expressed in laminae I and II, and part of laminae III in the spinal cord on the basis of co-expression with substance P, PAP and vGlut1 (Figure 1A). These data are consistent with earlier studies that showed co-expression of synaptophysin and Nav1.7 in lamina II of rat spinal cord (*14*). However, to our surprise, immuno-EM showed that the TAP-tagged Nav1.7 is present not only in presynaptic sites of central terminals of peripheral sensory neurons, but also in postsynaptic sites in dendrites in the spinal cord neurons as defined by ultrastructural criteria (Figure 1B) (*15, 16*). Quantitation of immunoreactive particles showed an association of 60% of immunoreactive Nav1.7 with postsynaptic sites where a third of immunoparticles were present on the membranes of dendritic shafts (Figure 1 C). Further immuno-fluorescence confocal microscopy was consistent with the finding that that the TAP-tagged Na_v_1.7 is expressed not only in the presynaptic terminals of sensory afferents but also in the postsynaptic terminals of interneurons in lamina II of the dorsal horn of the spinal cord (Figure S1).

**Figure 1.**
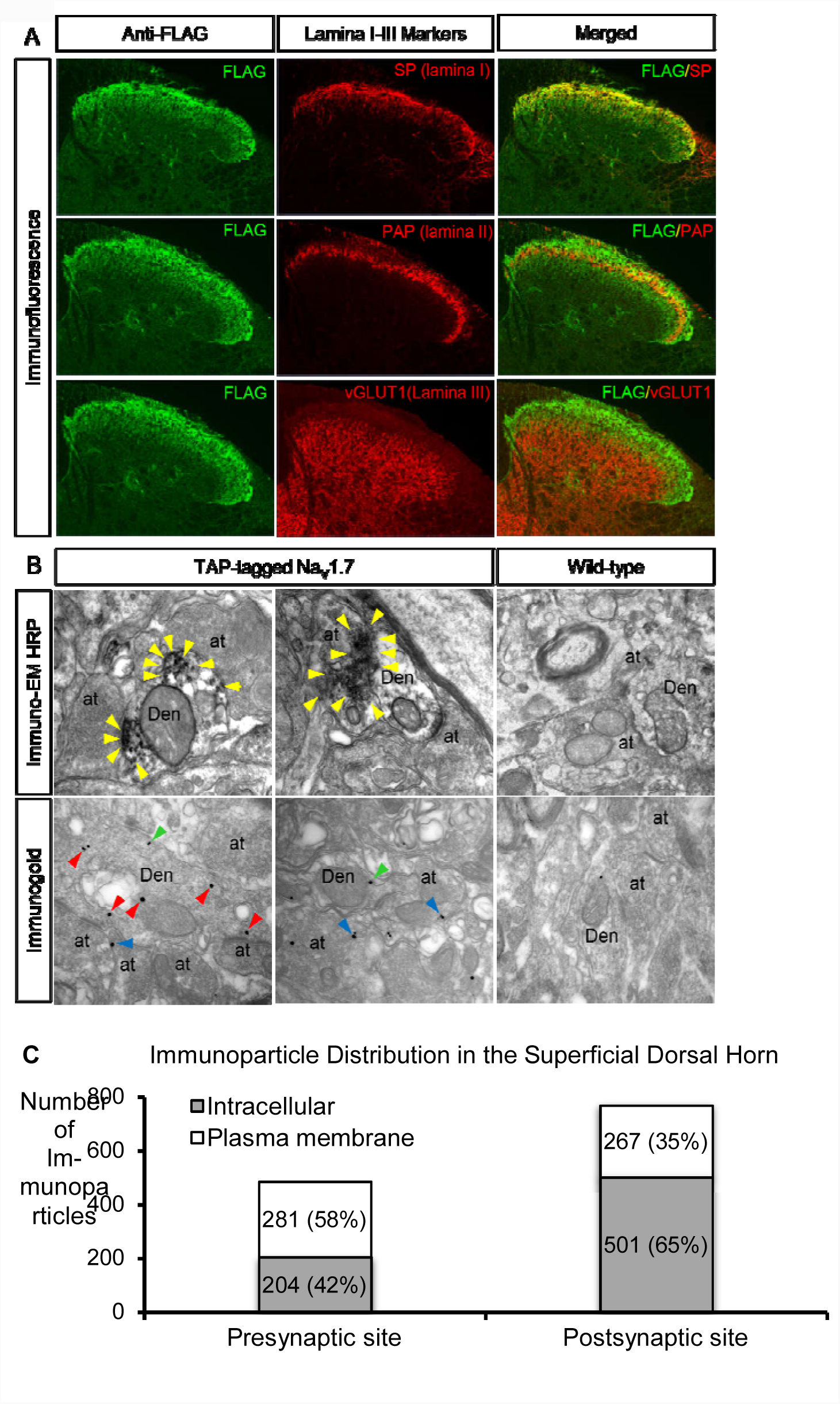
The distribution of Na_V_1.7 in the dorsal horn. A. the distribution of TAP-tagged Na_V_1.7 in the dorsal horn was detected using immunofluorescence with anti-FLAG antibody (left panels). The cross sections of spinal cord (lumbar 5) were co-stained with markers of superficial dorsal horn such as substance P (laminae I), PAP (laminae II) and vGLUT1 (Laminae III onwards) antibodies (middle panels in A). The right panels show the merged left to middle panels. The results show that TAP-tagged NaV1.7 mainly expresses in laminae I-II, and in the part of laminae III. Scale bar = 100 μm. **B**. the subcellular localisation of TAP-tagged Na_V_1.7 in the dorsal horn of the spinal cord were identified with immuno-electron microscopic techniques. Electron micrographs showing immunolabelling for FLAG in the dorsal horn of the spinal cord, were detected using the pre-embedding immunoperoxidase (top panels in B) and immunogold techniques (bottom panels in B). Using the pre-embedding immunoperoxidase method, peroxidase reaction end-product for FLAG in the TAP-tagged Na_V_1.7 mice was detected filling dendrites (Den) and dendritic spines of spinal cord neurons, as well at presynaptic sites filling axon terminals (at). In the wild-type mice (right panel in the top row), no peroxidase reaction end-product for FLAG was detected in the spinal cord. Using the pre-embedding immunogold method, FLAG immunoparticles present in the TAP-tagged Nav1.7 were mainly detected at intracellular sites (arrowheads in red), as well as along the plasma membrane (arrowheads in green) in dendritic shafts (Den) of spinal cord neurons. In addition, FLAG immunoparticles were observed along the extrasynaptic plasma membrane (arrow-heads in blue) of axon terminals (at). In the wild-type mice (right panel in lower row), a very low density of immunoparticles for FLAG, similar to background levels, was observed attached to mito-chondria in the spinal cord. Scale bars = 500 nm. **C**. the number of immunoparticles in the superficial dorsal horn of TAP-tagged Na_V_1.7 mice was counted. Out of 1253 immunoparticles, 768 particles were located at postsynaptic sites (61%) and 485 particles were located at presynaptic sites (39%). Along the 768 postsynaptic particles, 501 particles were located at intracellular sites (65%) and 267 particles were located along the plasma membrane (35%). Along the 485 presynaptic particles, 204 particles were located at intracellular sites (42%) and 281 particles were located along the plasma membrane (58%).

Does the dorsal horn neuron immunoreactive Nav1.7 originate from spinal cord mRNA transcripts? *In situ* hybridisation shows expression of Nav1.7 mRNA in a subset of motor neurons, but no transcripts in dorsal horn (*17*). These data therefore suggest that Nav1.7 protein or mRNA generated within sensory neurons is translocated to dorsal horn neurons. In order to explore any possible functional relevance of channel transfer, we examined the electrophysiological properties of dorsal horn lamina II neurons from mice where Nav1.7 is deleted only in sensory neurons using Cre-recombinase driven by the advillin promoter. Lamina II neurons exhibit different firing properties following somatic current injections and can be classified according to their firing pattern as tonic, burst, single spike or delay firing (*18*). We investigated whether there were differences in the relative frequency of each type of neuron between WT and DRG-specific Nav1.7KO mice. All four types of firing pattern were observed in both WT and Nav1.7KO animals as shown in the representative traces of Figure 2A-B. Interestingly, the prevalence of delay neurons was similar in WT (12/83) and Nav1.7KO (6/50) mice, but the percentage of tonic neurons in Nav1.7 KO mice was less than half that in WT (Nav1.7KO=18% vs. WT=40%) mice. The reduction in the proportion of tonic neurons in Nav1.7 KO mice was compensated by the prevalence of single spike (Nav1.7KO = 28% vs WT = 13%, Figure 2C-D) and to a lesser extent, bursting cells (42% vs 33%, Figure 2C-D.). The differences in neuronal populations in Nav1.7 KO compared to WT dorsal horn was statistically significant (p=0.0018, Chi-square test). Our data suggest that dorsal horn neurons from Nav1.7 peripheral neuron KO mice have reduced excitability. This was confirmed by the observation that the maximum number of spikes recorded during a 4 s maximal current pulse (sufficient to evoke firing at maximal frequency without spike inactivation) was reduced (Figure 2E, p=0.0288, two-sample t-test with Welch correction) with WT producing 51.7 ± 13.2 spikes (n=83 neurons) and Nav1.7 KO generating 26.9 ± 3.8 spikes (n=50 neurons). We also demonstrated that there is a significant increase in the AP threshold in Nav1.7 KO mice compared to WT neurons (Figure S2D, p=0.0483, two-sample t-test with Welch correction). As these events occur in mice where Nav1.7 is deleted only in sensory neurons, the deficits in dorsal horn neurons must result from a lack of translocated Nav1.7.

**Figure 2.**
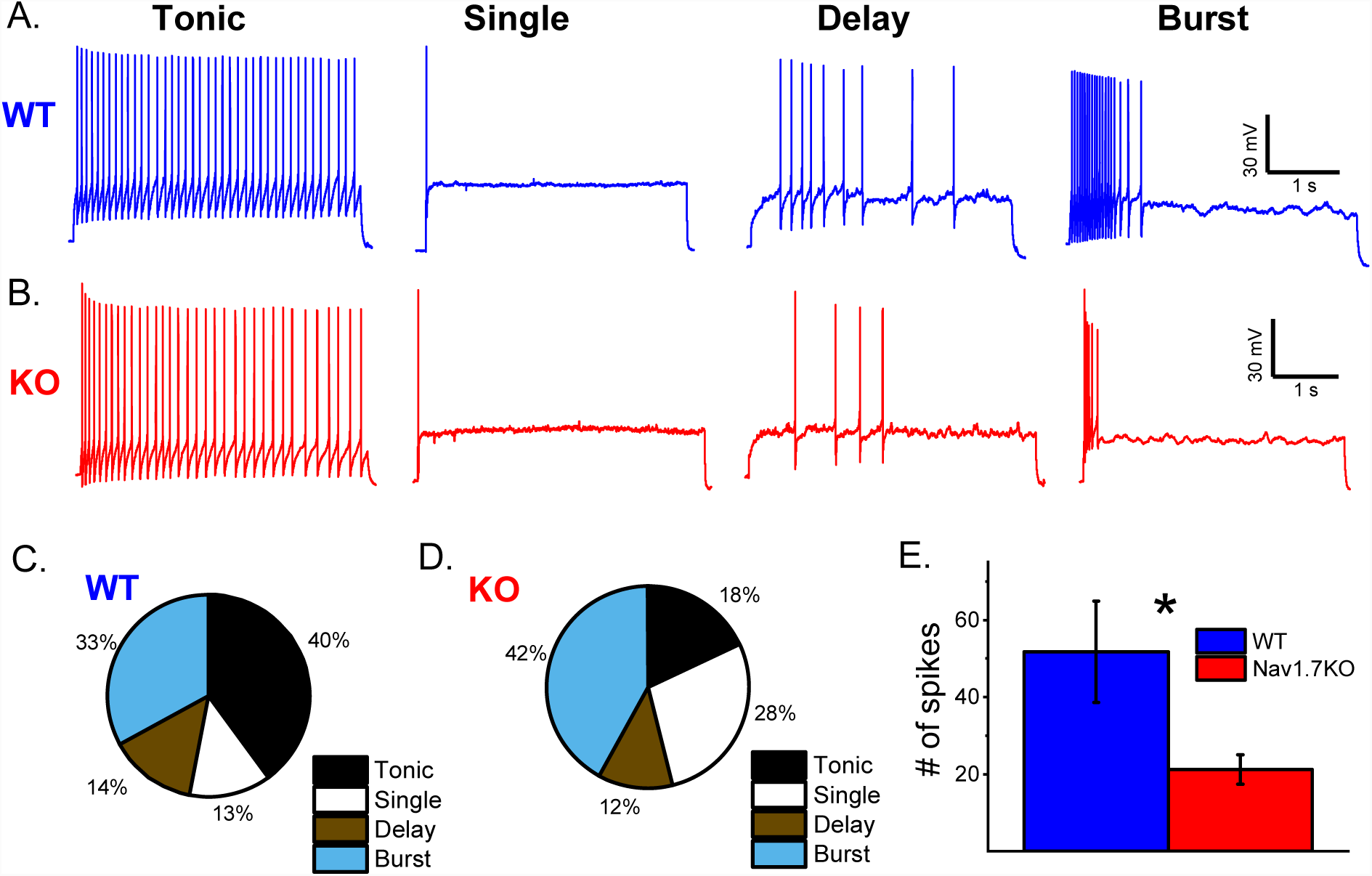
Electrophysiological properties of Lamina II neurons are altered in sensory neuron-specific Nav1.7 KO mice. Representative traces of the 4 main firing patterns of lamina II neurons recorded from *ex vivo* spinal cord slices from **A.** WT and **B.** Nav1.7 KO mice. Current injections are not shown for clarity. Current injections for recordings shown were: WT tonic = 80 pA, WT single = 200 pA, WT delay = 200 pA, WT burst = 60 pA, KO tonic = 30 pA, KO single = 100 pA, KO delay = 70 pA, KO burst = 50 pA. Percentage of each neuronal subpopulation in lamina II from **C.** WT (n=83 neurons) and **D.** Nav1.7 KO (n=50 neurons) mice. **E.** Maximum number of spikes elicited by current injection for each neuron compared between WT and Nav1.7 KO mice. The difference is statistically significant (*p=0.02875, two-sample t-test with Welch correction) with WT at 51.7 ± 13.2 spikes (n=83 neurons) and Nav1.7 KO at 21.2 ± 3.8 spikes (n=50 neurons).

We extended this genetic analysis by recording from Lamina II neurons in wild type mice to determine their firing properties before and after application of PF05089771 (henceforth PF771), a selective blocker of Nav1.7 channels (*19*). Application of the blocker changed the firing pattern in a minority of cells (6/24, Figure 3A, top panel). 3 burst firing neurons became single spiking neurons, 2 tonic firing neurons became burst firing and 1 tonic firing neuron became single spiking in the presence of PF771. In addition, in 10/24 neurons the rheobase significantly increased following application of PF771 (median 32.5 pA to 62.5 pA, p=0.00586, paired Wilcoxon signed rank test, Figure 3B). Consistently, in those cells in which PF771 increased rheobase, the median of the maximum number of spikes decreased from 26 to 6 spikes (p=0.0438, paired Wilcoxon signed rank test). In contrast, in the 14/24 cells in which rheobase was not significantly affected (median 25 to 22.5 pA, p=0.1, Figure S3B), the median number of spikes did not change significantly (median 33.5 and 36.5, p=0.358, paired Wilcoxon signed rank test, Figure S3C). In the neurons in which rheobase was increased, the voltage threshold was differentially affected as well: in the groups of cells responding to PF771 the median voltage threshold was −37.6 mV in control and increased to −34 mV (p = 0.00805, paired Wilcoxon signed rank test), while it did not change for the non-responding cells (median −37.4 to −38 mV, p = 0.286, Figure S3D).

**Figure 3.**
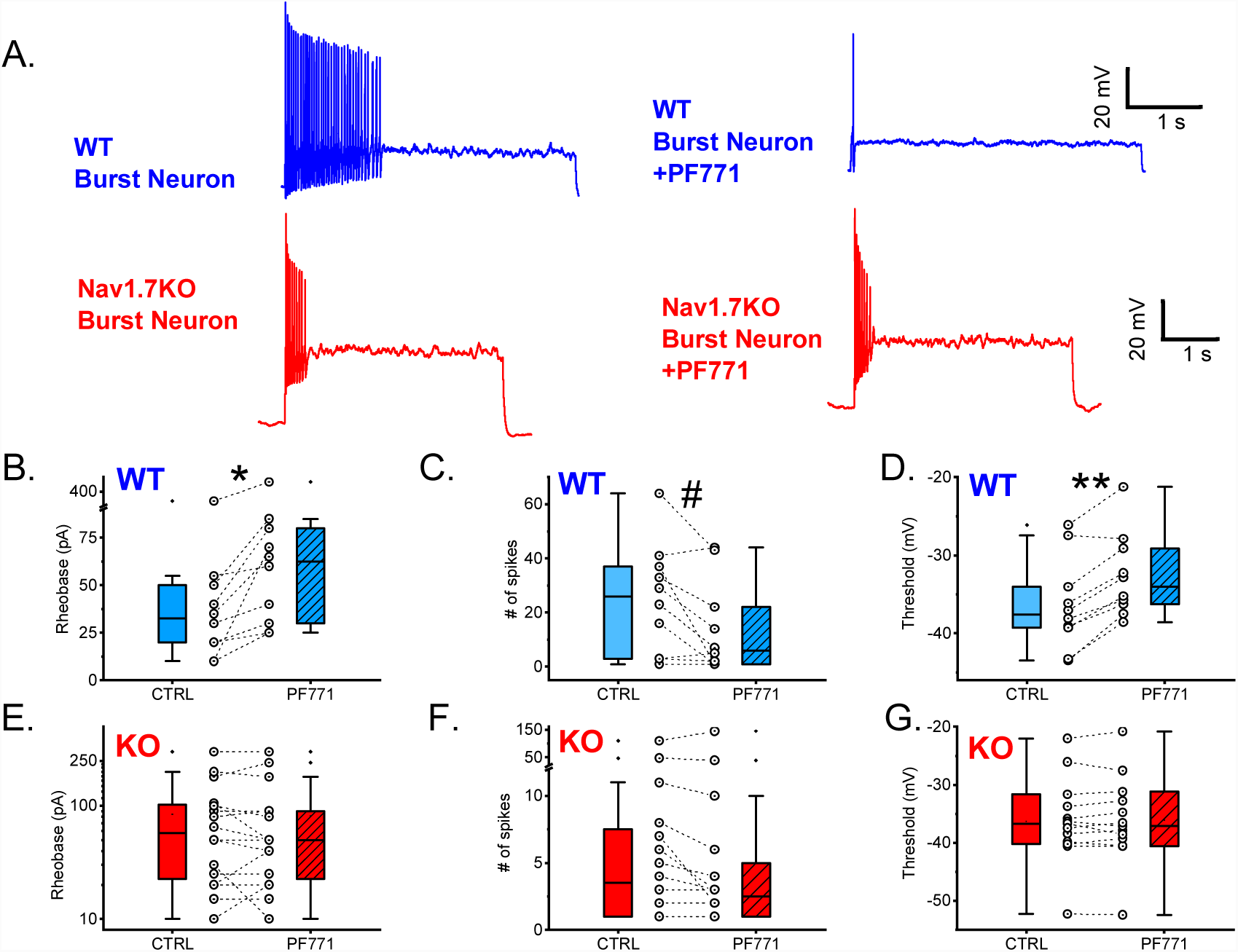
Effect of specific Nav1.7 blocker PF-05089771 on lamina II neurons from wild-type and Nav1.7 KO mice. Two populations of WT neurons were identified: neurons that displayed an increase in rheobase and neurons that showed little or no change in rheobase in the presence of PF771 (see Figure S2). **A.** Representative WT burst firing neuron displaying an effect of PF771 on firing. **B.** Representative Nav1.7 KO burst firing neuron displaying no effect of PF771 on firing. Current injections for traces were as follows: WT burst = 30 pA, and Nav1.7 KO burst = 50 pA. Current injections were identical before and after drug. **B.** WT Neurons were split on the basis of rheobase change in the presence of PF771. Paired WT neurons displayed an increase in rheobase with PF771 (hatched bar) vs. control (open bar, *p=0.00586, paired Wilcoxon signed rank test) **C.** Maximum # of spikes (at the same current injection before and after drug) was significantly reduced with PF771 vs. control (#p=0.04383, paired Wilcoxon signed rank test) in WT neurons showing an increase in rheobase only. **D.** Threshold was significantly increased with PF771 vs. control (**p=0.00178, paired Wilcoxon signed rank test) in WT neurons showing an increase in rheobase only. There were no major population differences in terms of response to PF771 identified in Nav1.7 KO mice dorsal horn neurons. There were no significant changes in **E.** rheobase **F.** # of spikes or **G.** threshold of Nav1.7 KO dorsal horn neurons in the presence of PF771 vs. control.

It is worth noting that the block by PF771 at this concentration is only effective after channels have been inactivated by a prolonged depolarization (cells that exhibited a change in input resistance of 10% or more following this depolarisation were discarded from further analysis, see Methods). Since any Nav1.7 channels located on primary afferents would not have been inactivated and would be insensitive to PF771, the effects observed are specific to postsynaptic Nav1.7 on dorsal horn neurons. These data confirm that functional Nav1.7 channels are expressed on the postsynaptic membrane of a subset of Lamina II neurons and their activation contributes to neuronal excitability.

In order to confirm the specificity of the action of PF771, we repeated the same experiments on Nav1.7 KO mice, measuring the electrophysiological properties of n=16 Lamina II neurons. In Nav1.7 KO mice, there was no significant change in rheobase (median 57.5 pA to 50 pA, p=0.504), maximum number of spikes (median 3.5 to 2.5, p=0.181) or threshold (−36.7 mV to −37.1 mV, p=0.514) in the presence of PF771 (Figure 3E-G). This indicates that knockout of Nav1.7 in presynaptic sensory neurons results in a functional deficit post-synaptically in the superficial dorsal horn.

Our functional studies demonstrate that the excitability, threshold and firing pattern of a subset of dorsal horn neurons can be ascribed to the presence of sensory neuron-derived Nav1.7. These findings have significance for analgesic drug development, and help to explain the failure of peripherally restricted Nav1.7 antagonists to be effective analgesics (*9*). Nav1.7 loss, apart from effects on sensory neuron excitability, increases endogenous opioid drive, inhibits neurotransmitter release and, as we have shown here, diminishes dorsal horn excitability. The relative importance of these regulatory steps in Nav1.7 loss-of-function analgesia is uncertain. Nonetheless, our findings demonstrate intercellular transport of Nav1.7, a functional role for the channel in recipient cells, and a pattern of Nav1.7 expression that differs from that predicted by transcriptomic studies. Nav1.7 antagonists have yet to produce as profound analgesia as loss of the encoding gene itself, probably because complete channel block seems to be required to induce opioid-peptide expression (*7*). These results support the view that targeted ablation of *scn9a* in sensory neurons may be a more productive approach to analgesia than the development of small molecule antagonists of Nav1.7.

What is the mechanism of trans-neuronal transfer of Nav1.7? Evidence for protein transfer between cells has been obtained in the immune system (trogocytosis) (*20*), with transcription factors (*21*), through trans-synaptic transfer and via exosome release(*22*). The trans-neuronal transfer of synuclein has been linked to neurodegenerative disease (*23*). Recent studies also demonstrate trans-synaptic transfer of RNA via the activity-dependent immediate early retrotransposon capsid-like ARC protein that is linked to synaptic plasticity, memory formation and neuropsychiatric disease (*24*). It seems that circuitry within the brain involves not only electrical signalling but also activity-dependent transfer of RNA and proteins. This is the first example of peripherally encoded proteins exerting effects on central nervous system function. However, the mechanism of Nav1.7 transfer remains to be established.

## Acknowledgements

We gratefully acknowledge support from Wellcome 10154/Z/13/Z, 200183/Z/15/Z Versus Arthritis and EU Horizon 2020. This work was supported by grants from the Junta de Comunidades de Castilla-La Mancha (PPII-2014-005-P) to R.L. and by a Medical Research Council grant (MR/R011494/1) to MB. We thank David Attwell and Fan Wang for helpful criticism, and Andrew Todd and Janka Erdos for assistance with immunohistochemistry experiments.

