## Supplementary data for "Sensory neuron-derived Na_v_1.7 contributes to dorsal horn neuron excitability"

**SUPPLEMENTARY MATERIAL – Alles et al., 2019**

**Materials and Methods**

***Animals***

Both female and male mice aged 4-8 weeks were kept on a 12-h light/dark cycle and provided with food and water *ad libitum*. Conditional Nav1.7 KO mice were generated by crossing floxed (SCN9A) Nav1.7 mice with Advillin-Cre mice^1^. Nav1.7^TAP^ knock‐in mice were generated as described previously^2^. All experiments were performed with approval from the United Kingdom Home Office according to guidelines set by personal and project licenses, as well as guidelines of the Committee for Research and Ethical Issues of IASP.

**Immunofluorescence**
The perfusion and staining were performed as previous described (Zhao et al, 2010). Briefly, after deep anaesthesia with pentobarbitone sodium solution (80 mg/kg, Pentoject, 57-33-0, Animal-care) by intraperitoneal injection (e.g. 20μl/25g mouse), animals were transcardially perfused with 10 ml heparinized saline (0.9% w/v NaCl; 10 units/ml heparin, Wockhardt UK Ltd.) followed by 25 ml of fresh prepared 4% (w/v) paraformaldehyde in 0.1 M phosphate buffer (PB) pH 7.4 (containing 15% saturated picric acid and 0.1% glutaraldehyde for Immuno-EM). After perfusion, the spinal cords were removed, post-fixed in the same fixative solution at 4°C overnight, and then cryopro-tected in 30% w/v sucrose containing 0.02% sodium azide in 0.1 M PB overnight. Subsequently, the sections were cut at 60 µm thickness on a vibratome (VT1000S, Leica), and then incubated in 50% ethanol for 30 minutes, blocked in a blocking buffer (1x PBS containing 0.3% Triton X-100 containing 5% donkey serum) at the room temperature for 1 hour, and then incubated in primary antibodies diluted in the blocking buffer at 4°C overnight. After 3 washed in PBS, the sections were incubated in the secondary antibodies at room temperature in dark for 2 hours. Later, the sections were rinsed 3 times in PBS and then placed onto Superfrost plus (Ref J1800AMNZ, Thermo Scien-tific) slides, mounted in an Antifade Mounting Medium (H-1000, Vector), covered with coverslip and seal with nail varnish. Finally, the sections were scanned with a Zeiss LSM 710 204 confocal mi-croscope. The primary antibodies are anti-FLAG (1:100, F1804 Sigma); anti-Substance P (1:200, OBT06435, Oxford Biotec); anti-PAP (1:1000, AAF23171 Aves Lab Inc.) and anti-vGLUT1 (1:5000, AB5905, Chemicon), anti-Homer (1:1000, Frontier Institute, AB_2631104), anti-vGLUT2 (1:5000, ab2251, Millipore). The secondary antibodies are donkey anti-mouse A488 (1:1000, Cat No Molecular Probes), donkey anti-chicken Rhodamine Red (1:1000, Cat. No Molecular Probes),donkey anti-rat-A647(1:1000, Cat.No, Molecular Probes), donkey anti-guinea pig Pacific Blue (1:1000, Cat. No Molecular Probes) and donkey anti-goat Rhodamine red (1:1000, Cat. No Molecular Probes)

***Immunohistochemistry for electron microscopy (Immuno-EM)***

Immunohistochemical reactions for electron microscopy were carried out using the pre-embedding immunoperoxidase and immunogold methods described previously^3^. Briefly, 10 week-old TAP-tagged Na_V_1.7^KI/KI^ mice (n=3) and wild-type littermate control mice (n=3) were used for Immuno-EM. After deeply anesthetized, the cross sections from lumber 4-5 of spinal cords were prepared as above (Immunofluorescence). Subsequently, the free-floating sections were blocked, incubated with a monoclonal anti-FLAG (3.5 μg ⁄ mL, F1804, Sigma) antibody in blocking buffer (tris-buffered saline (TBS) containing 1% (v/v) normal goat serum (NGS)). they were incubated in biotinylated goat anti-mouse IgG (Vector) or in goat anti-mouse IgG coupled to 1.4 nm gold (Nanoprobes Inc., Stony Brook, NY, USA) diluted in TBS containing 1% NGS. For immunoperoxidase, the sections were then transferred to an avidin-biotinylated peroxidase mixture (ABC kit, Vector) diluted as 1:100 for 2 hours at room temperature. Peroxidase enzyme activity was revealed by using a 3,3´-diaminobenzidine tetrahydrochloride solution (DAB; 0.05% in TB, pH 7.4), to which 0.01% H_2_O_2_ was added. For immunogold, the sections were post-fixed in 1% (v/v) glutaraldehyde and washed in double-distilled water, followed by silver enhancement of the gold particles with an HQ Silver kit (Nanoprobes Inc.). All sections were then treated with osmium tetraoxide (1% in 0.1 m phosphate buffer), block-stained with uranyl acetate, dehydrated in graded series of ethanol and flat-embedded on glass slides in Durcupan (Fluka) resin. Regions of interest were cut at 70-90 nm on an ultramicrotome (Reichert Ultracut E, Leica, Austria) and collected on single slot pioloform-coated copper grids. Staining was performed on drops of 1% aqueous uranyl acetate followed by Reynolds’s lead citrate. Ultrastructural analyses were performed in a Jeol-1010 electron microscope.

***Spinal cord preparation***

Spinal cord preparations were obtained from male or female mice, between 30 and 60 days old, from either wild-type C75Bl/6 (WT) or conditional Nav1.7 knockout (Nav1.7 KO). Animals were anesthetized via intraperitoneal injection of a ketamine/xylaxine mix (80 mg/kg and 10 mg/kg respectively) and decapitated. The spinal cord was dissected in ice cold aCSF of the following composition: (in mM) 113 NaCl, 3 KCl, 25 NaHCO3, 1 NaH2PO4, 2 CaCl2, 2 MgCl2, and 11 D-glucose and the same solution was used for subsequent electrophysiological recordings. Once dissected free from the vertebral column, the spinal cord was carefully cleaned from connective tissues and dorsal roots were cut at approximately 2 mmm length. The spinal cord was then glued to an agar block and glued to the slicing chamber of a HM 650V vibratome (Microm, ThermoFisher Scientific, UK). The solution used for slicing contained (in mM) 130 K-gluconate, 15 KCl, 0.05 EGTA, 20 HEPES, 25 D-glucose, 3 kynurenic acid, 2 Na-Pyruvate, 3 Myo-Inositol, 1 Na-L-Ascorbate, and pH 7.4 with NaOH^4^. Slices were incubated for 40 minutes at 35 degrees and then allowed to equilibrate at room temperature for further 30 minutes before starting the recordings.

***Electrophysiology***

Current and voltage clamp recordings were performed using either a Molecular Devices Multiclamp 700B (Scientifica, UK) or an ELC-03X amplifier (NPI electronics, Germany). Signals were filtered at 5KHz, acquired at 50 KHz using a Molecular Devices 1440A A/D converter (Scientifica, UK) and recorded using Clampex 10 software (Molecular Devices, Scientifica, UK). Electrodes were pulled with a Flaming-Brown puller (P1000, Sutter Instruments, USA) from borosilicate thick glass (GC150F, Harvard Apparatus, UK). The resistance of the electrodes, following fire polishing of the tip, ranged between 3 and 5 MΩ. Bridge balance was applied to all recordings. Intracellular solution contained (in mM) 125 K-gluconate, 6 KCl, 10 HEPES, 0.1 EGTA, 2 Mg-ATP, pH 7.3 with KOH, and osmolarity of 290–310 mOsm. Cells were included for further analysis if their input resistance throughout the duration of the experiment was stable (within 10% of its initial value) and if their spikes (evoked by a current step) peaked beyond 5 mV.

Cells were targeted in the inner and outer Lamina II and visualized through an Eclipse E600FN Nikon microscope (Nikon, Japan) equipped with infrared differential interference contrast (IR-DIC) connected to a digital camera (Nikon, DS-Qi1Mc). In most experiments a dorsal root was stimulated via a section electrode connected to an isolated current stimulator (DS3, Digitimer, UK). The stimulation intensity was fixed at 5×threshold for evoking the low threshold response in the recorded cell (typically 50-100 µA).

Cells were held in current clamp mode and their resistance and capacitance were measured from the voltage response to a brief (20 ms) current step (10-20 pA). Rheobase was defined as the minimum current necessary to elicit a spike. Cells were classified into tonic (firing continuously throughout a current step), single spike (firing only once at the beginning of the current step), delayed (firing during the course of the current step) and bursting (firing more than one spike at the start of the current step).

PF05089771 was bath applied through perfusion at a concentration of 100 nM. It has been shown that at our chosen concentration PF05089771 has no blocking effect on Nav1.7 unless the channels are inactivated^5^. Indeed, in our experimental conditions, the response to current injection was never affected by the presence of the blocker. The drug effects were observed only after Nav1.7 channels were inactivated by holding the cell at depolarized potential (0 mV) for 30 s. Cells that exhibited a change in input resistance of 10% or more following this depolarisation were discarded from further analysis. Cells were defined as ‘responsive’ to the channel blocker if their rheobase increased by at least 10% in the presence of PF05089771.

***Drugs***

All drugs were obtained from Sigma-Aldrich unless stated otherwise.

***Data analysis***

For electrophysiological data, the spike kinetic parameters (spike width, height, threshold and maximum rate of rise and descent) were measured using custom written software (Bayesian Quantal Analysis suite, freely downloadable at <http://sourceforge.net/projects/pyclamp/>^6^).

All other statistical and data analysis was performed with Origin Lab 2018, GraphPad Prism 7.0, Microsoft Excel or Zeiss Zen Blue microscopy software.

**Supplementary Figures**

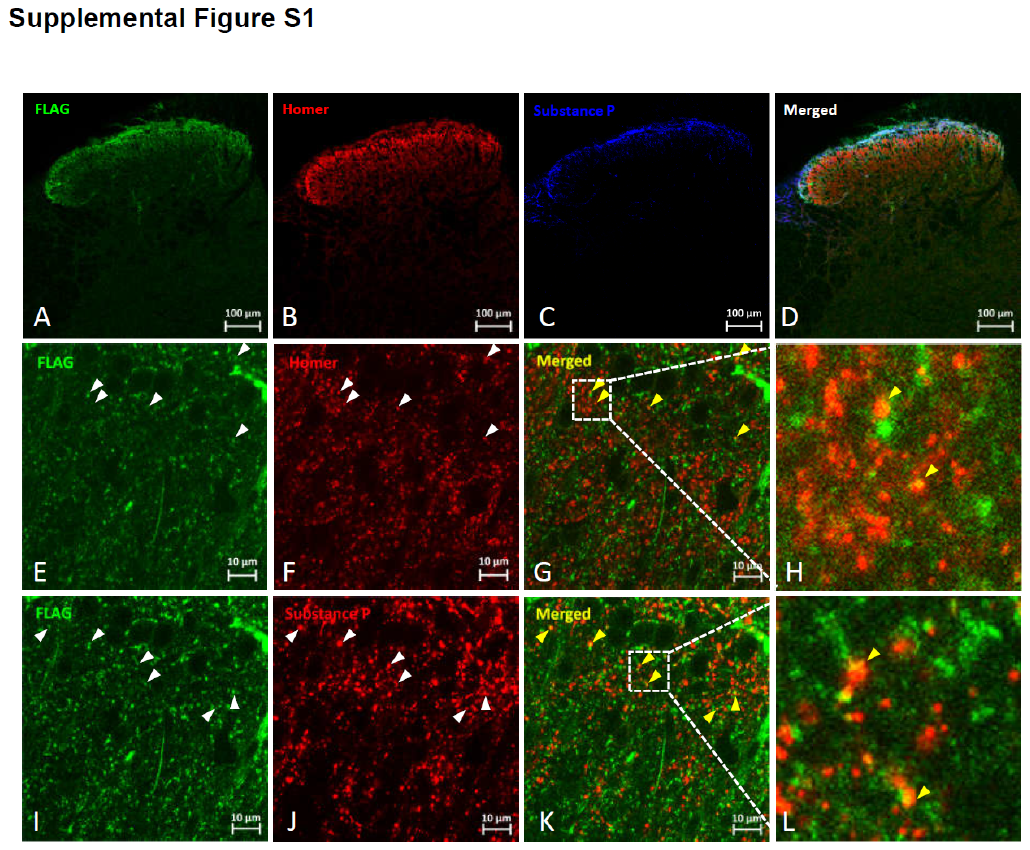

**Supplemental Figure S1. The distribution and co-localisation of Na_V_1.7 in the dorsal horn.** The distribution and co-localisation of Na_V_1.7 in the dorsal horn were also detected using immunofluorescence and confocal microscopy with both presynaptic marker Substance P (SP) and postsynaptic marker Homer, especially for excitatory synapses of interneurons in dorsal horn. (A-D) Low magnification (20x) views show that the immunoreactivity of FLAG (TAP-tagged Na_V_1.7) is present in Lamina I, II and part of III compared to Homer mainly in Lamina II and III, and Substance P in Lamina I. Higher magnification (63x) views of parts of Lamina II (E-H) and Lamina I (I-L) show that the immunostainings are in the form of small puncta that are scattered throughout the neuropil (K-I and O-P). The yellow dots pointed by headarrows (in yellow) in panel G and H, and in panel K and L indicate the co-localisation of Na_V_1.7-Homer and Na_V_1.7-SP in superficial Lamina II and I, respectively. Scale bars = 100 μm (A-D), and 10 μm (E-G and I-K).

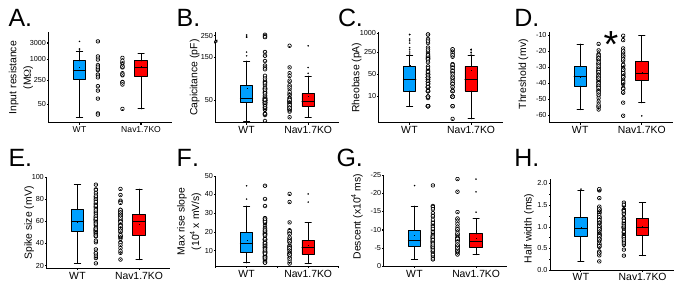

**Supplemental Figure S2. Intrinsic properties of WT and Nav1.7 KO superficial dorsal horn neurons. A.** Input resistance (in MΩ) **B.** capacitance (pF) **C.** rheobase (pA) **D.** threshold (mV)**. E.** spike size (mV) **F.** max rise slope (x10^4^ mV/ms) and **G.** descent (x10^4^ ms) and **H.** AP half-width (ms) of WT (light blue, n=83 neurons) and Nav1.7 KO dorsal horn neurons (red, n=50 neurons). Threshold was the only property that was significantly increased in Nav1.7 KO dorsal horn neurons compared to WT dorsal horn neurons (*p=0.04834, two-sample t-test with Welch correction). Interestingly, it has been shown that there is a decrease in AP threshold in nociceptors derived from chronic pain patients with inherited erythromelalgia (IEM) harbouring a gain-of-function mutation in the Nav1.7 channel, which agrees with our data^7^. Input resistance, capacitance, rheobase, spike size, max rise slope, descent and half width were not significantly changed (p>0.05, two-sample t-test with Welch correction) in Nav1.7 KO compared to WT dorsal horn neurons.

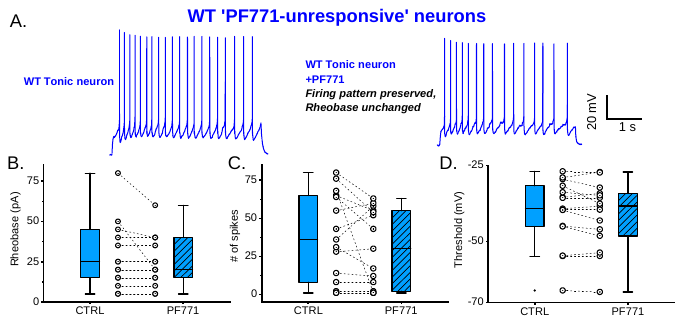

**Supplemental Figure S3. ‘PF771-unresponsive’ WT superficial dorsal horn neurons**. Two populations of WT neurons were identified: neurons that displayed an increase in rheobase and neurons that showed little or no change in rheobase in the presence of PF771. **A.** Representative WT tonic firing neuron displaying no effect of PF771 on firing pattern or rheobase. Current injection was 60 pA both before and after drug. There were no significant changes in **B.** rheobase **C.** # of spikes or **D.** threshold of these neurons in the presence of PF771.
